# Slice-sampled Bayesian PRF mapping

**DOI:** 10.1101/093724

**Authors:** Silvan C. Quax, Thomas C. van Koppen, Pasi Jylänki, Serge O. Dumoulin, Marcel A.J. van Gerven

## Abstract

Functional magnetic resonance imaging (FMRI) allows to non-invasively measure human brain activity at the millimeter scale. As such, it is widely used in computational neuroimaging studies that aim to build models to predict stimulus-induced neural responses in visual cortex. A popular method is population receptive field (PRF) mapping, which is able to characterize responses to a large range of stimuli. For each voxel, the PRF method estimates the best fitting receptive field properties (such as location and size in the visual field) using a coarse–to–fine approach which minimizes, but not eliminates, the risk of returning a local minimum. Here, we provide a Bayesian approach to the PRF method based on the slice sampler. Using this approach, we provide estimates of receptive field properties while at the same time being able to quantify their uncertainty. We test the performance of conventional and Bayesian approaches on simulated and empirical data.

## 1 Introduction

In visual neuroscience, the investigation of receptive field properties has been one of the major advancements in understanding the visual system. Conventionally, the term receptive field has been used to describe the region of visual space that, when illuminated, would trigger a response in a retinal neuron [1]. In modern usage, the term receptive field can be used in recording sites along the entire visual pathway, and has been expanded to include stimulus features such as orientation, motion, direction, and color. Early electrophysiological studies in monkeys have found an orderly structure in which input from the retina is mapped onto the early visual cortex [2, 3, 4]. This structure respects topology, in a way that neighboring locations in the visual field are nearby in the primary visual cortex (V1). Furthermore, it was found that the entire visual field is represented multiple times throughout the visual cortex, corresponding to different visual areas [5, 6]. Receptive field properties (i.e., location of the receptive field) provide a way to construct the preferred positions in the visual field of each recording site, from which visual field maps can be constructed. Furthermore, by creating a model for the receptive field, one can predict neural responses to different stimuli. Recording techniques that are used in investigating human receptive field properties often take the responses of many neurons. In creating receptive field models at this scale, one refers to the model as a population receptive field (PRF) model as it measures the pooled response from several million neurons [7].

In humans, functional magnetic resonance imaging (FMRI) provides an excellent instrument to measure brain activity as a response to visual stimuli, and is often used in visual field mapping. Conventional visual field maps are obtained by phase encoded methods that estimates a single location in the visual field that, when stimulated, evokes the largest blood oxygenated level dependent (BOLD) response in the visual cortex [8, 6]. However, several studies quantified a range of possible locations that effectively activated a voxel in the early visual cortex [9, 10]. This formed the foundation for a framework to non-invasively estimate PRF properties for FMRI that was introduced by Dumoulin and Wandell [11]. The PRF method provides a model for the receptive field of a voxel in the visual cortex and a way to estimate receptive field properties (i.e. location, size). By modeling receptive fields, the method by Dumoulin and Wandell provides a way to compare modeled properties obtained by different recording instruments, as they are expressed in units of visual space (i.e. degree visual angle) as opposed to instrument-specific units (e.g., BOLD, FMRI; voltage, LFP). Additionally, this method is able to estimate PRF properties using a wide range of visual stimuli, as opposed to the widely used moving wedge and contracting ring stimuli in phase-encoded retinotopic mapping.

The standard PRF approach uses a binary representation for the stimulus aperture and combines this with a model for the population receptive field. Often this receptive field model is taken to be a symmetric two-dimensional Gaussian, but many extensions have already been made to allow for more complex models that include surround suppression or use asymmetric shapes [12]. The PRF model is estimated on a voxel-by-voxel basis using a two-stage approach which consists of a coarse stage and a fine stage, where the best fitting model parameters are fitted by minimizing the residual sum of squares (RSS). In the coarse stage a grid of roughly 100.000 plausible combinations of PRF parameters (location: *x*_0_,*y*_0_ and width: *ϕ*), are used to create predictions. The coarse stage is done on a smoothed cortical surface, as this imposes a spatial correlation between neighboring voxels. The fine stage uses voxels from the coarse stage to fine-tune the model parameters on a non-smoothed surface. This method increases processing time, but maximizes the likelihood of finding a global minimum. For each voxel, the method effectively provides the best set of PRF parameters from the data.

Here, we develop a Bayesian inference approach for the PRF method which provides us with underlying distributions for each model parameter, and gives us information on how certain we are of each parameter’s estimate. The underlying distributions add a new way of assessing data. For instance, we can investigate how this distribution behaves under different task conditions. It has been shown that receptive fields are not rigid over time, but can change due to attention effects or task demands [13]. This new method enables us to quantify how variable our underlying receptive field is by the uncertainty of the posterior estimate. Additionally, we can express our beliefs in the distribution of model parameters by defining (informative) priors based on knowledge about the distribution of receptive field densities and sizes [14, 11]. By incorporating priors, we make the method more robust when using less data, or under noisy observations. To test our Bayesian PRF framework, we use simulated data to test for performance compared to the PRF method. We evaluate our model on fMRI data to retrieve empirical PRF maps.

## 2 Methods and Materials

### 2.1 Population receptive field model

Here we describe the PRF method as a starting point to define our Bayesian model, for a more detailed explanation of the PRF method we refer the reader to the paper by Dumoulin and Wandell [11].

The PRF model assumes that the voxel response to a stimulus can be accurately described in terms of a receptive field parameterized by a set of parameters *θ*. In case of the two-dimensional Gaussian PRF model, which we describe here, these parameters are center location m = (*m_x_, m_y_*), and width (or standard deviation of the Gaussian) *ϕ*. The goal in PRF modeling is to fit the best set of parameters by minimizing the residual sum of squares between prediction and data.

The stimulus at time *t* is described by a function *s*(x, *t*) that marks the stimulus intensity in the visual field at position x = (*x, y*) (in degrees of visual angle).

Let *gθ* (*x*) denote a response function parameterized by *θ* characterizing how a voxel responds to a point stimulus at location x. The neuronal population response *r*(*t*) is given by the product between the stimulus and the receptive field

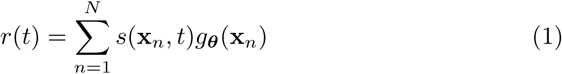

when summing over all *N* possible locations (pixels). We can more conveniently write the responses at all time points *t_k_* with 1 ≤ *k* ≤ *T*, in matrix notation as

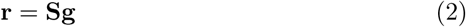

where **S** is a *T* × *N* matrix such that *s_kn_* = *s*(*x_n_*,*t_k_*) and g is a *N* × 1 vector such that *gn* = *gθ* (x*_n_*).

The neuronal response is indirectly measured at times *u_k_* with 1 ≤ *k* ≤ *M*. Within a neuroimaging setting, these measurements depend on a convolution of the neural response with a hemodynamic response function (HRF) *h*(*t*). The predicted response is thus given by:

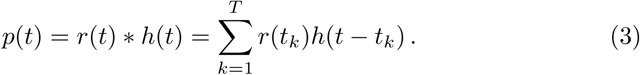

This can be written in matrix notation for all *M* measurement times as

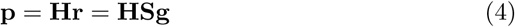

where

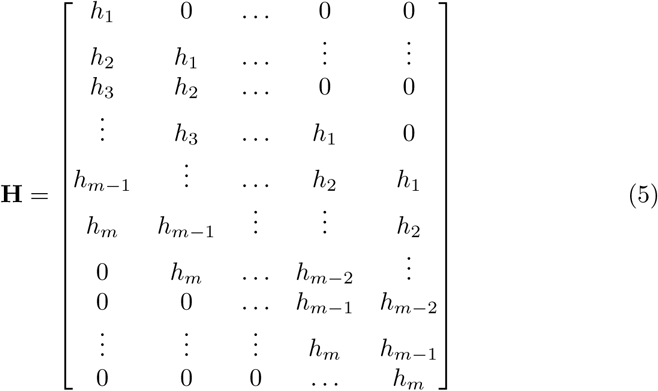

is the *M* × *T* convolution (Toeplitz) matrix.

We express the measured BOLD timeseries in one voxel as

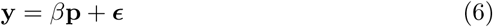

where *β* is a scaling factor and *∈_t_* ∼ 𝒩(0,*σ*^2^) is Gaussian white (measurement) noise.

The model is completed by choosing a particular functional form for the response function which characterizes a neuron’s receptive field. Here, for convenience, we assume that it is given by a Gaussian function

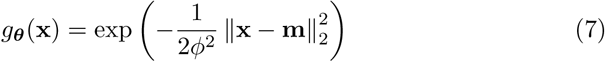

with parameters *θ* = (**m**, *ϕ*^2^), where **m** = (*m_x_*, *m_y_*) is the center of the receptive field and *ϕ^2^* is the size of the receptive field. Using these definitions we can write down the likelihood function

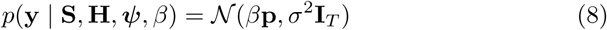

where we used *ψ* = (*θ*, *σ*^2^).

### 2.2 Bayesian estimation of population receptive fields

In a Bayesian setting, the goal is to compute the posterior density for the parameters of interest *ψ* given the stimuli **S**, the HRF **H**, and the observed data y. In order to write down this posterior, we require the likelihood (8) as well as suitable priors on the parameters. These terms may in turn depend on suitably chosen hyper-parameters (given by *ξ*). In the appendix we define the required priors, which can be uninformed or chosen based on physiological knowledge.

For our purposes, we are not interested in *β*, as it is simply a scaling factor that contains no information for our receptive field model. Therefore, we interpret the scaling factor as a nuisance parameter, and we can integrate it out after we approximate our posterior distribution (see below). Using our apriori assumptions for the scaling factor *β* and noise *∈* we note that our noise and scaling factor are both drawn from a Gaussian. Using Equation (6) and invoking properties of multivariate Gaussian distributions we can write the likelihood function (8) as:

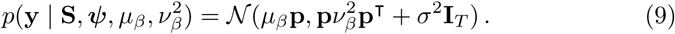

### 2.2.1 Approximate inference

To find the distributions for our model parameters, we would like to compute the posterior:

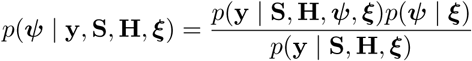

which follows from Bayes’ rule. The left hand side of this equation, the posterior, is the probability of our parameters after observing our data. The right hand side numerator is given by a product of the *likelihood,* which is the probability of observing the data given the parameters, and the parameter priors, which expresses our beliefs in the model parameters before observing. The right hand side denominator is the *marginal likelihood* and is a normalizing constant that involves a multidimensional integral, which is often costly to compute.

We approximate the posterior using a Markov chain Monte Carlo (MCMC) approach, which draws samples *ψ^t^* from the posterior, where *ψ^t^* depends on the previously drawn sample *ψ*^*t*-1^. MCMC does not require evaluation of the normalizing constant *p*(**y** | **S, H, ξ**) = ∫ *p*(**y**, *ψ* | **S, H, ξ**) *dψ*. If we take enough samples using our MCMC method, we approximate the distribution well enough to avoid computing the marginal likelihood. For our priors we assume independence between parameters. This allows us to write the prior as a product of individual beliefs in our parameters:

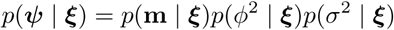

For our MCMC method, we use the slice sampler [15] to generate draws from the posterior. This method allows us to approximate the posterior without having to solve the integral, even if we add more parameters. To prevent numerical issues, and to increase the speed of drawing samples, sampling proceeds in the log domain (see Appendix A for a derivation of the required log joint PDF).

## 2.3 Simulation

To validate the Bayesian PRF model, we compare the model as well as the original PRF fitting procedure by Dumoulin and Wandell [11] against ground truth by using simulations. The simulations were generated using the Gaussian PRF model defined before, which both analysis techniques assume as the ground truth model in this case. Validation of both models was done by testing the efficiency and accuracy with which the parameters of the ground truth model were recovered.

We test this model on data simulated using the standard moving bar stimuli and an HRF that was obtained from data from one subject in a prior experiment. The bar stimuli consisted of a bar (of width 1.56 degree visual angle) spanning over the entire stimulus window (radius 6.25) moving in four orientations (0°, 45°, 90°, and 135°) for two different motion directions, giving a total of 8 different bar passes. Additionally, 4 mean luminance (blank) periods were inserted, giving a total of 12 periods (8 bar passes/4 blanks), each lasting 20 frames of 1.5 seconds. For each voxel, we generate two-dimensional symmetric Gaussian RFs using some **m** and *ϕ*. To generate the data we multiplied the known RFs with stimulus aperture at each time and convolved them with a fixed HRF. We chose the combinations of RF parameters (centers,between −1.00 and 1.00 degree visual angle; spread, between 0.25 and 1.50 degree visual angle) so that 99.98 percent of the receptive fields would stay within the stimulus window (-6.25 to 6.25 degree visual angle) to minimize issues that might arise due to boundary effects [16]. We chose the scaling factor *β* to be 1, and generated 1280 voxels with added random white noise with increasing amplitude *σ_J_* to create the training set. The noise amplitude corresponded with a signal-to-noise (SNR) ratio as given by:

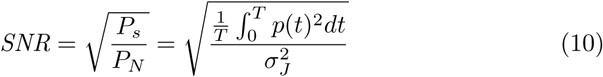

We repeated this process with the same parameters to create the test set. For both the Bayesian PRF model and the conventional PRF we assumed a known HRF and provided the HRF that was used in generating the data. Figure 1 shows the simulated timeseries of a voxel with an RF centered at **m** = [0.17,0.57] with *ϕ* = 1.24.

We tested for convergence using the convergence statistic described in Appendix B. For each voxel we ran four chains to test for convergence. As starting points for our slice sampler we choose random initial values for each chain for **m** (between −8 and 8 deg), *ϕ* (between 0 and 8 deg), *σ*^2^ (between 0 and 2 times the variance in the timeseries).

**Figure 1:**
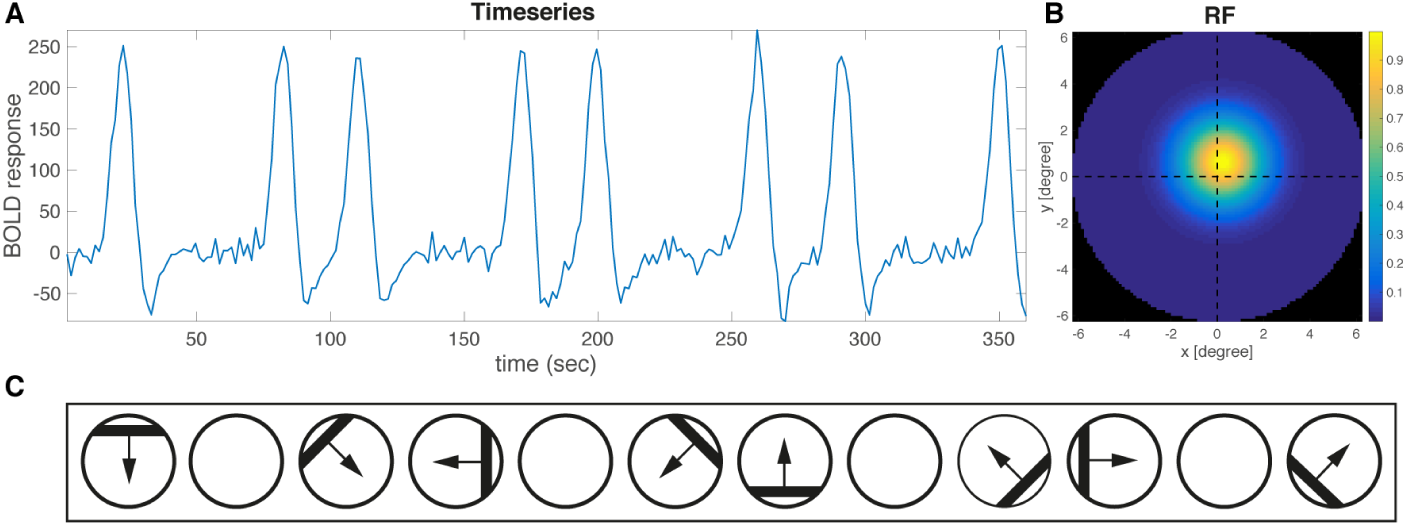
Simulated BOLD response of a voxel centered at **m** = [0.17,0.57] with *ϕ* = 1.24 (**A**). Coresponding receptive field (**B**). The stimulus aperture sequence included 8 bar passes and 4 blanks (**C**).

## 2.4 Empirical data

To see how our model performs on real neuroimaging data we tested our model on a previously obtained FMRI data set [12]. This data set consisted of ∼160000 voxels in the posterior half of the brain of a single subject. The stimuli used were the same moving bars as in the simulations. This resulted in 240 time points of data per voxel. The FMRI data provided ROIs that were drawn on a gray/white segmented (inflated) cortical surface from estimates obtained with the PRF method. Additionally, the data provided a HRF that was obtained during the study. For the Bayesian model, we had to follow the preprocessing steps (e.g., detrending, percent-bold-conversion) used in the PRF method as described in [11]. Furthermore, we ran the Bayesian model with non-informative priors on location and PRF size. Since the data consisted of 12 scans of the same stimuli, we could divide the data into a training set and a test set. This enabled us to validate our estimated models predictions against the test set. We ran our estimation analysis with an uninformed (flat) prior as well as with a weakly informed prior. Initial tests with a strong prior turned out to bias the estimation to strongly. The weakly informed prior consisted of a normal distribution 𝒩(*μ* = 0, *σ* = 8) over the center of the receptive field, a lognormal distribution LogNormal(*μ* = 3, *σ* = 1.8) over the receptive field size and a lognormal distribution LogNormal(*μ* = 3, *σ* = 1.3) on the noise.

## 2.5 Choosing priors

In our simulation study we have used uninformative priors. For real data, we also looked at the influence of an informative prior. These priors are based on previous literature concerning general relationships of receptive fields in the human brain.

**Prior: location** For the location parameter **m**, we assume independent normal distributions for the *x* and *y* directions:

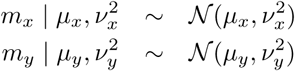

In choosing the parameters *μ_x_*, 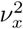, *μ_y_*, and 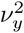 we use literature on the cortical magnification factor (CMF). The CMF is a measure of neuronal organization in the visual cortex and is defined as the cortical surface distance between two points with visual field position spaced 1° apart [2]. We can use the CMF to construct our prior on location, as the CMF is a measure of how many neurons are ‘dedicated’ to a part in the visual field, assuming a constant neuron density within different areas of visual cortex [17]. A higher CMF means that more neurons are selective to that part in the visual field. In other words, we can think of the CMF as corresponding to the PRF density over the visual field. Several studies have investigated the properties of the CMF in humans using FMRI [14, 18].

For the primary visual cortex (V1), studies show a linear relationship between inverse cortical magnification factor *M*^-1^ and eccentricity, as described by fitting the following equation:

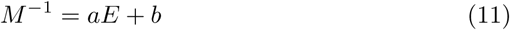

where *E* is eccentricity, and *a* and *b* are fitting parameters. Duncan & Boynton [14] found a linear relationship between the inverse magnification factor and eccentricity:

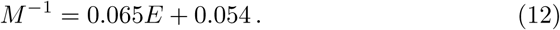

Using this linear relation for *M*^-1^ we can identify a suitable prior for location parameter **m**, using *M* (i.e., the PRF density) as our measure for the probability density function. We use this to estimate the variance *ν*^2^ for our prior on location according to a zero mean normal distribution.

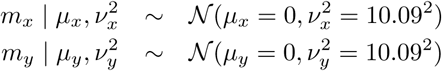

**Prior: Receptive field size** Similarly, we would like to construct a prior for the receptive field size *ϕ*^2^. As we expect more smaller receptive fields due to over representation of foveal locations, we suggest a candidate function being a log-normal distribution given by:

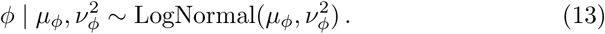

To obtain values for the hyperparameters *μ_ϕ_* and 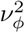 for the prior on location, we look at two known linear relationships. The first is the linear relationship between the CMF and eccentricity as described above. The second is the relationship between PRF size and eccentricity. Linear fits to data for visual field maps in V1-V3 have been found [11]. Using this data we can obtain a linear relationship for V1, taken from the average of the three areas:

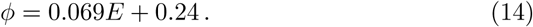

We can now combine this relationship with the relationship of the CMF with eccentricity by the PRF size.

If we now use the linear relationship between *ϕ* and eccentricity we can fit our log-normal distribution. We obtain the following prior:

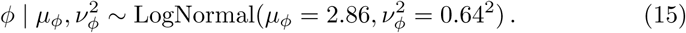

By using the M-scaling as a basis for the priors on location and *ϕ*, we do violate the conditional independence requirement for our priors, that enables the factorization of our posterior. Additionally, we should note that both the CMF-eccentricity and the PRF size-eccentricity relationships vary for each visual area. Ideally, one would have to obtain different relationships for these areas to construct priors for each visual area. Additionally, one would have to define the areas by obtaining initial PRF estimates by running a non-informative approach.

## 3 Results

### 3.1 Simulation

To test how our model behaves we ran several simulations. Since our method is able to quantify the uncertainty in the estimation of our parameters it is interesting to see how this uncertainty changes with noise in the simulated data. Here we show the sampled marginal distributions after 600 iterations (taking only the last 400 iterations) for voxels under three different noise levels. Additionally, we show the model predictions for both the Bayesian and the conventional PRF method. Our potential scale reduction factor converged to 1 within 600 samples for the data set.

In Figure 2, we show a sampled distribution for a voxel with a high signal-to-noise ratio (SNR=9.35). For this voxel, the Bayesian estimates lie closer to the ground truth values than the conventional estimates for both location and PRF size. The distributions are highly confined in parameter space (as shown by the interval range on the horizontal axis), showing that we are certain of our estimates. The predicted timeseries from both the conventional and the Bayesian method are similar for these noise levels.

**Figure 2:**
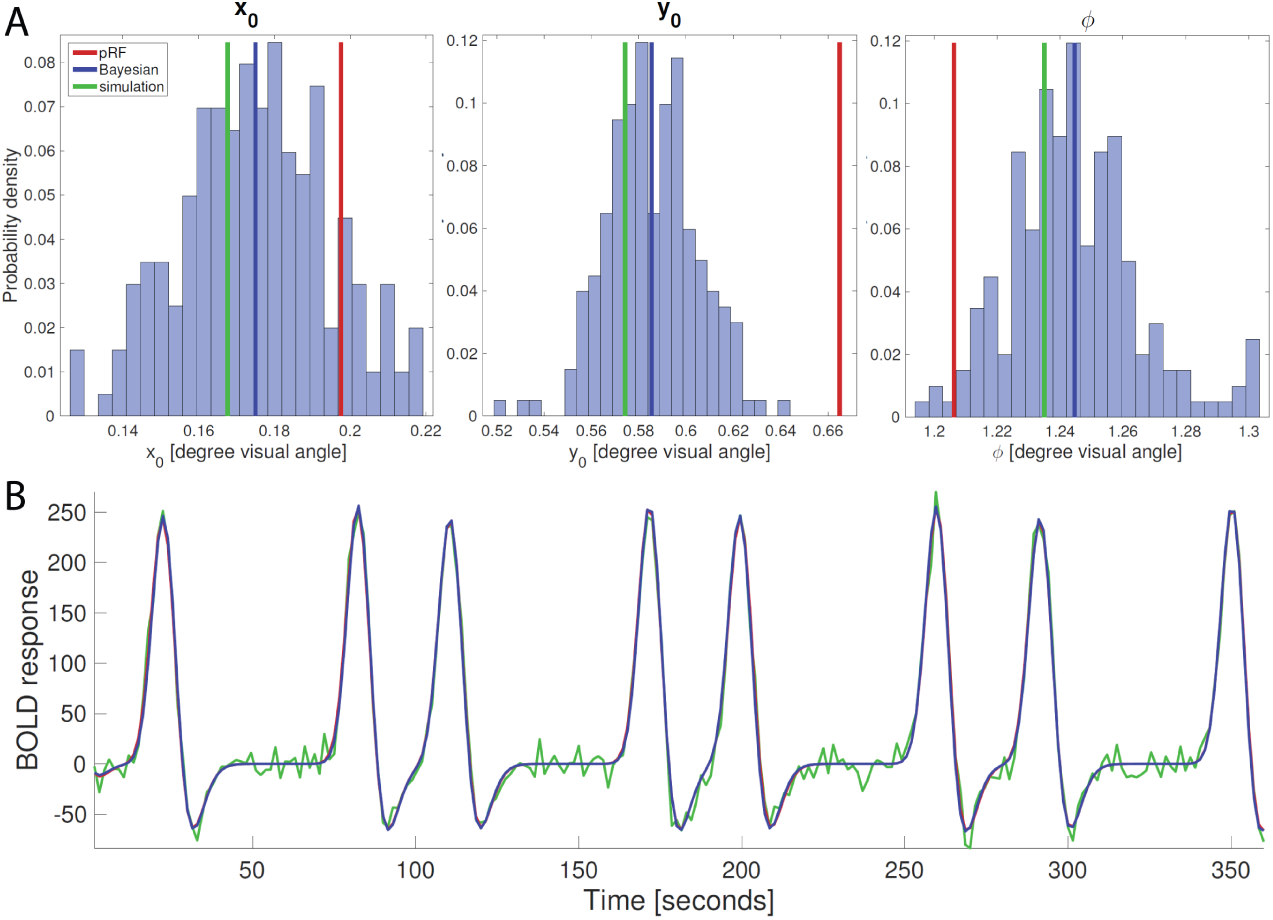
The sampled probability densities (of which the last 400 are shown) of slice sampling for a single voxel under low noise conditions (SNR=9.35) are shown in panel A. Green indicates simulation (true) values, red indicates PRF estimates, and blue indicates median estimates using the Bayesian method. For this voxel, the Bayesian estimates lie closer to the simulation values for both location and PRF size. Panel B shows the resulting predicted timeseries, where the simulated data is shown in green. No clear differences in predictions between the Bayesian (blue) and the PRF method (red) can be seen, as they overlap.

Figure 3 shows the results for a voxel with a more realistic signal-to-noise ratio (SNR=1.71). Here we see that the distributions are wider than the low noise condition, showing we are less certain of our estimates. For this voxel, we see that the *y*_0_ estimate from the PRF method is closer to ground truth than the Bayesian method. The estimates for *x*_0_ and *ϕ* are slightly closer to ground truth for the Bayesian method. Again, the predicted time series for both methods are very similar.

**Figure 3:**
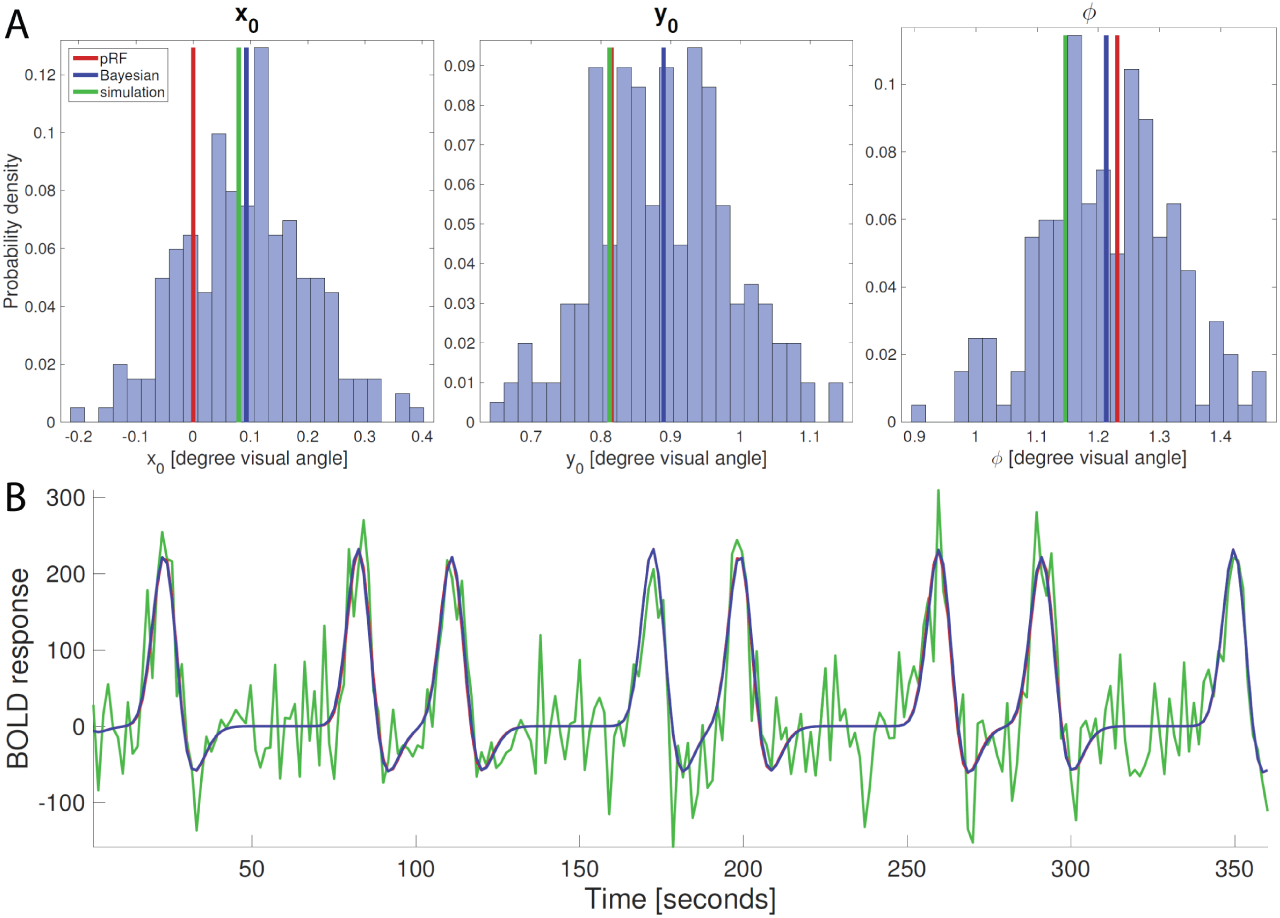
The sampled probability densities for a voxel under realistic noise conditions (SNR=1.71). For each parameter, we see the probability densities span a larger portion of parameter space, showing that we are less certain of our estimates compared to the low noise condition. The predicted timeseries show no clear differences in predictions between the Bayesian and the PRF method.

Figure 4 shows the results for a voxel with a very low signal-to-noise ratio (SNR=0.17) to illustrate what happens under highly noisy conditions. Here we can see that the distributions span over the entire slice sampler window, indicating that we are completely uncertain of our parameter estimates. No information is conveyed by both the conventional and the Bayesian model. We should note that the relative proximity of the ground truth values and the PRF estimates for the location parameter *x*_0_ and *y*_0_ do not signify a better fit. Instead, they are the result of our location parameter ground truth values chosen at center locations (between −1 and 1 degree visual angle), and the conventional method defaulting near these values when no suitable fit can be found in the grid fid stage.

**Figure 4:**
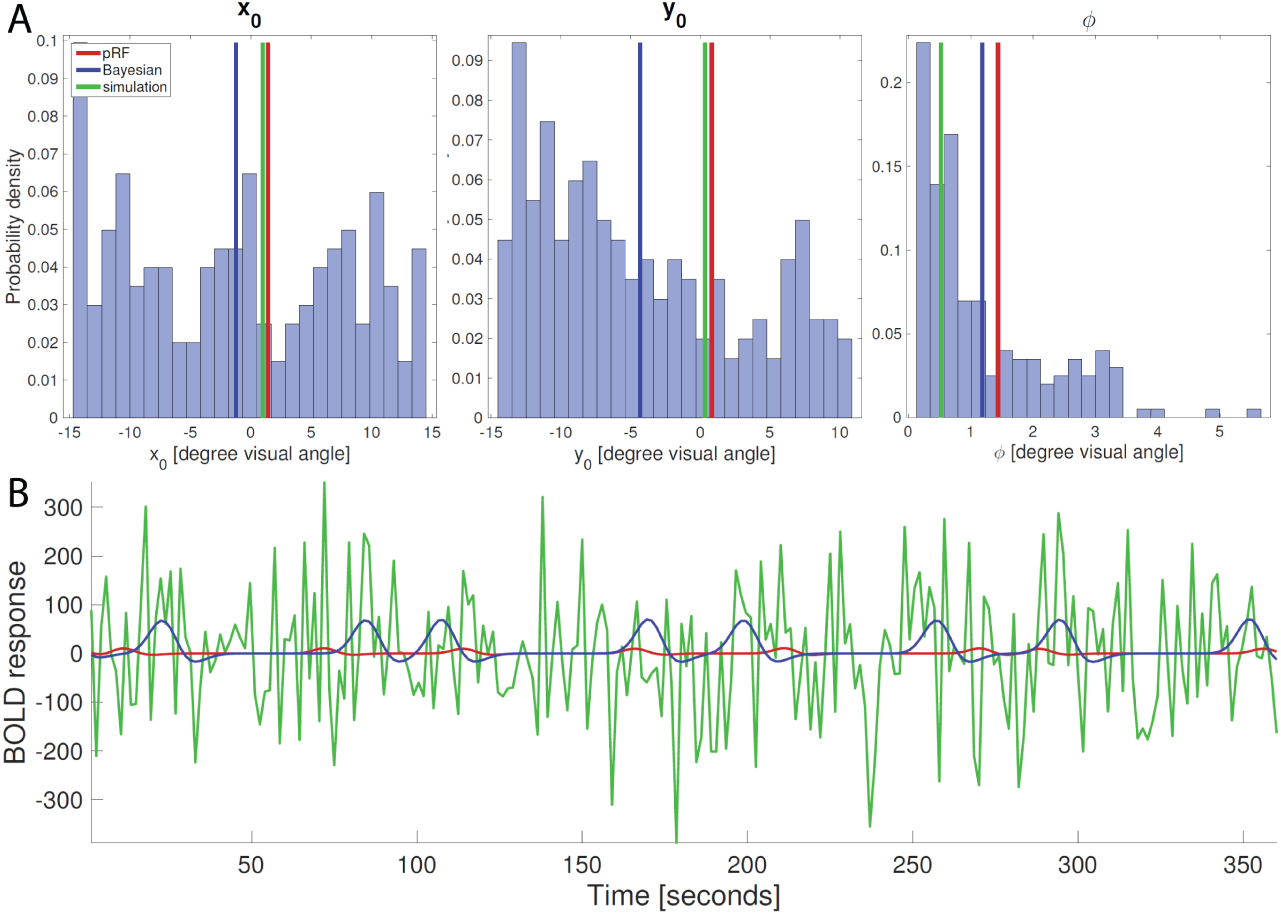
The sampled probability densities for a voxel under high noise conditions (SNR=0.17) that was taken from a different data set. Under these conditions, the distributions span over the entire slice sampler window. The predicted time series clearly differ, but both convey no information on the data.

Figure 5 shows the performance of the conventional and the Bayesian approach on the test set. Predictions that explained less than .15 (the lower threshold in many PRF experiments) of the variance in the test set were omitted, resulting in N=1074 for the Bayesian method and N=1068 for the conventional method. The first panel (A) shows the performance in estimating the RF center position by the conventional (red) and the Bayesian method (blue). Here, the Bayesian model performed better in retrieving ground truth location estimates for all noise levels in the test set. Panel B shows the absolute error in RF size with increasing noise levels. Similarly, the Bayesian model outperformed the conventional method in retrieving PRF size for all noise levels. Panel C shows the explained variance of the predictions on the test set, with an upper bound of .15 explained variance. The explained variance did not show significant differences between the Bayesian and the conventional method.

**Figure 5:**
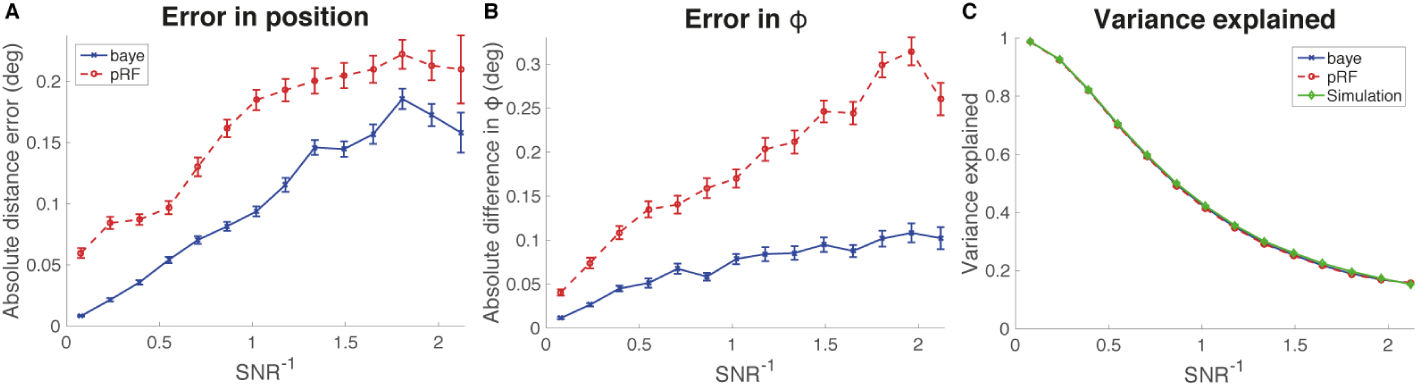
Performance of the PRF and BPRF methods on the test set (N=1280 voxels, after thresholding N=1074 for BPRF; N=1068 for PRF). Panel A shows the error in absolute distance between simulated RF centers and RF centers obtained by the Bayesian (blue) and PRF (red) with increasing noise levels (inverse of the signal-to-noise ratio (SNR). Panel B shows the absolute error in RF size (*ϕ*). Panel C shows the explained variance on the test set by the prediction with RF properties estimated in the training set. The upper bound for the *SNR^-1^* corresponds with 0.15 explained variance, we chose this value as this is the usual lower threshold to include voxels during in-vivo PRF experiments.

### 3.2 Empirical data

To validate our analysis technique we used previously obtained data (see Methods) [12]. We checked whether we can obtain PRF maps similar to other techniques. For illustration purposes, we show a voxel in V1 that contained a signal with a reasonable SNR. Figure 6 shows the resulting distributions and prediction for both the median Bayesian estimate (blue), and the conventional estimate (red). For this voxel, both methods provide similar estimates for the location parameters, and differ slightly in PRF size estimates.

**Figure 6:**
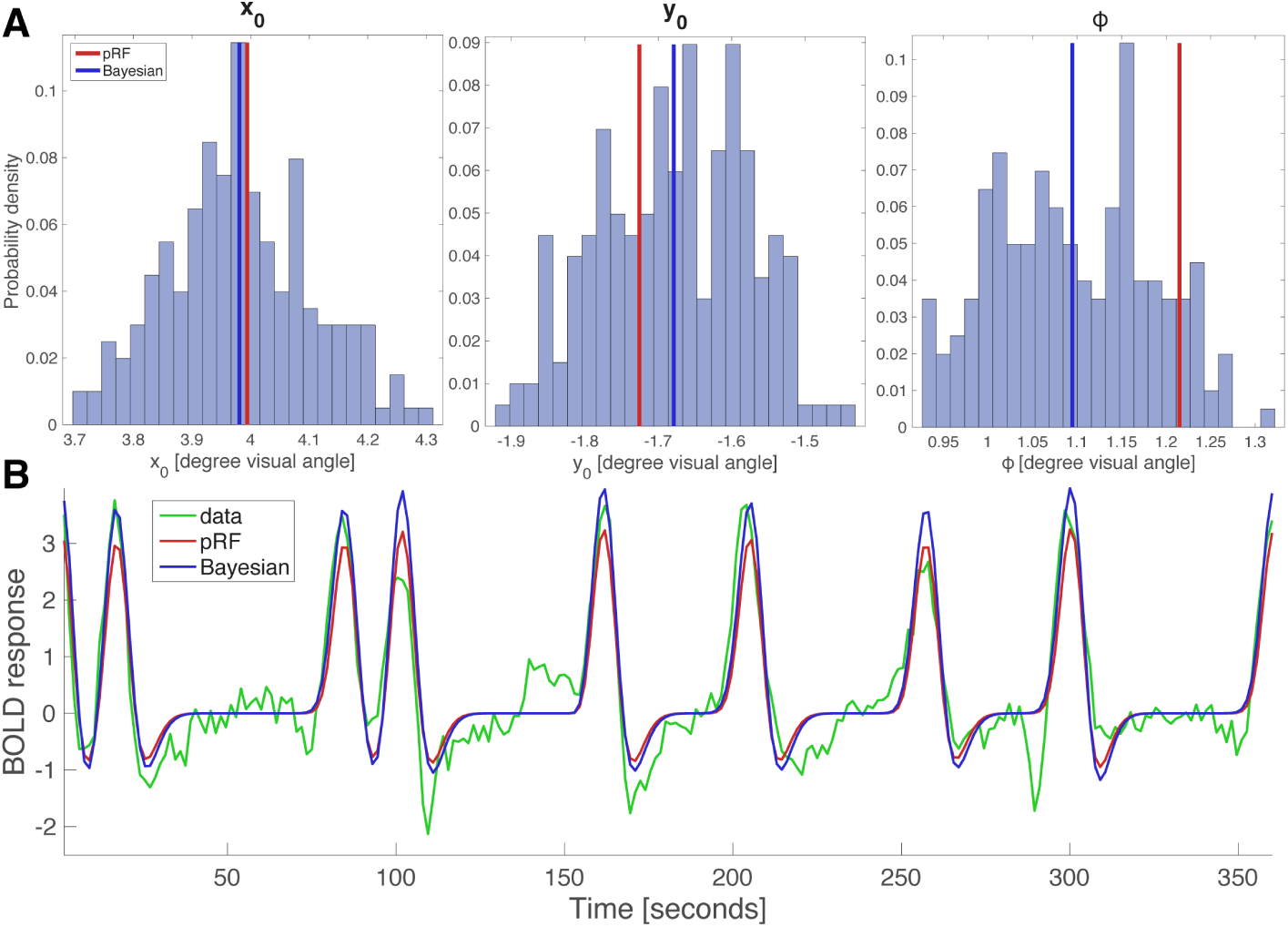
Example run on a voxel from real data obtained from an experiment. A shows the sampled distributions after 600 samples for *x*_0_, *y*_0_ and *ϕ*. Red shows PRF estimates and blue shows median Bayesian estimates. B shows the predicted response, here the data (converted to percent BOLD and detrended) is shown in green.

To see whether our model retrieves PRF maps similar to [11] we ran our analysis on all 164912 voxels. Initially we used an uninformed prior to see how our model performed without any prior knowledge. Our sampling method takes on average ∼ 30 seconds to estimate the parameters of a single voxel, though this can vary greatly depending on noise level and how well our model fits the data. Luckily every voxel is estimated independently so we can run multiple sampling chains in parallel. Using 50 nodes on a computing cluster this resulted in *∼*6 hours needed to estimate PRF parameters for one subject.

The results of our PRF map estimation are shown in Figure 7. The eccentricity, receptive field size and polar angle are shown for voxels with an explained variance above 0.15. Because we now also have a measure of the uncertainty of our estimations we can make uncertainty maps for the different estimated parameters for all voxels (Figure 8).

**Figure 7:**
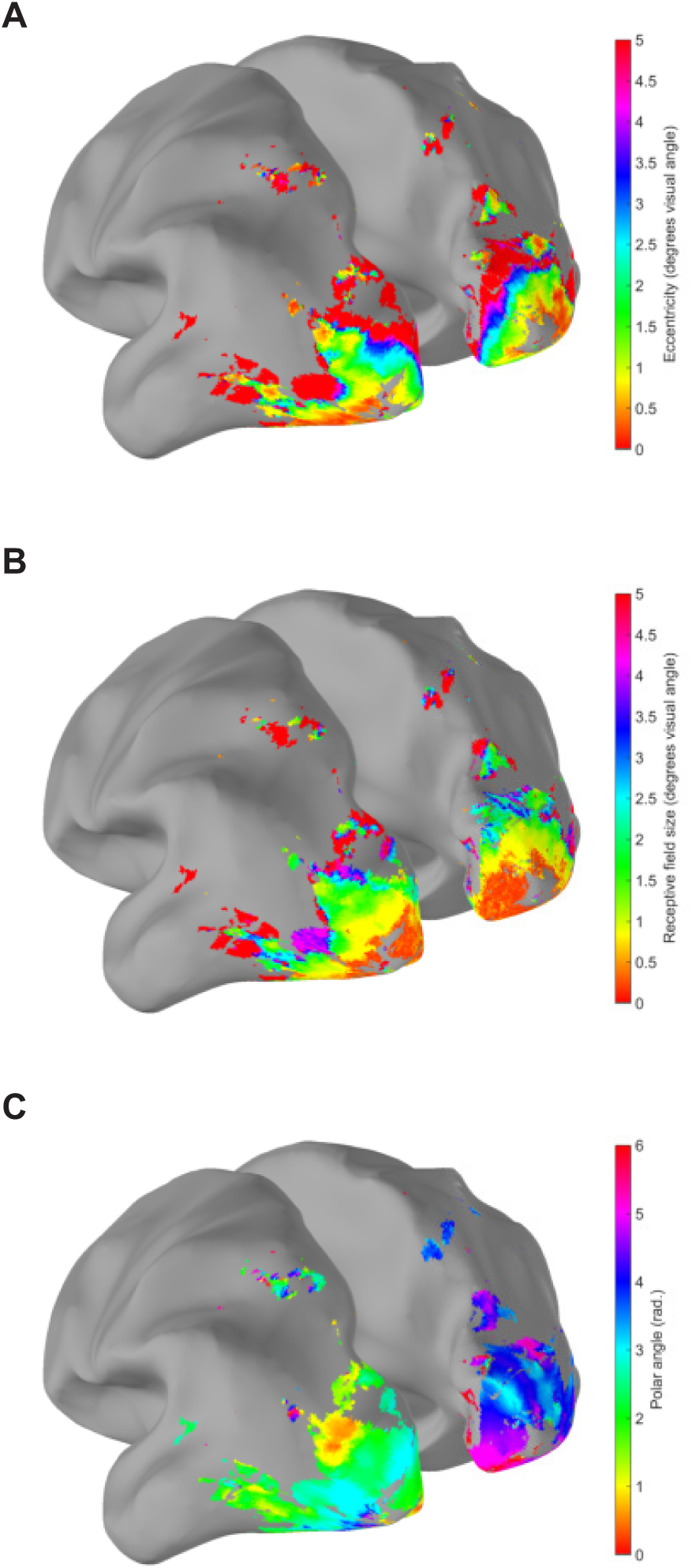
Maps of PRFs in the cortex as estimated by our Bayesian approach. From top to bottom: eccentricity, PRF size, polar angle.

**Figure 8:**
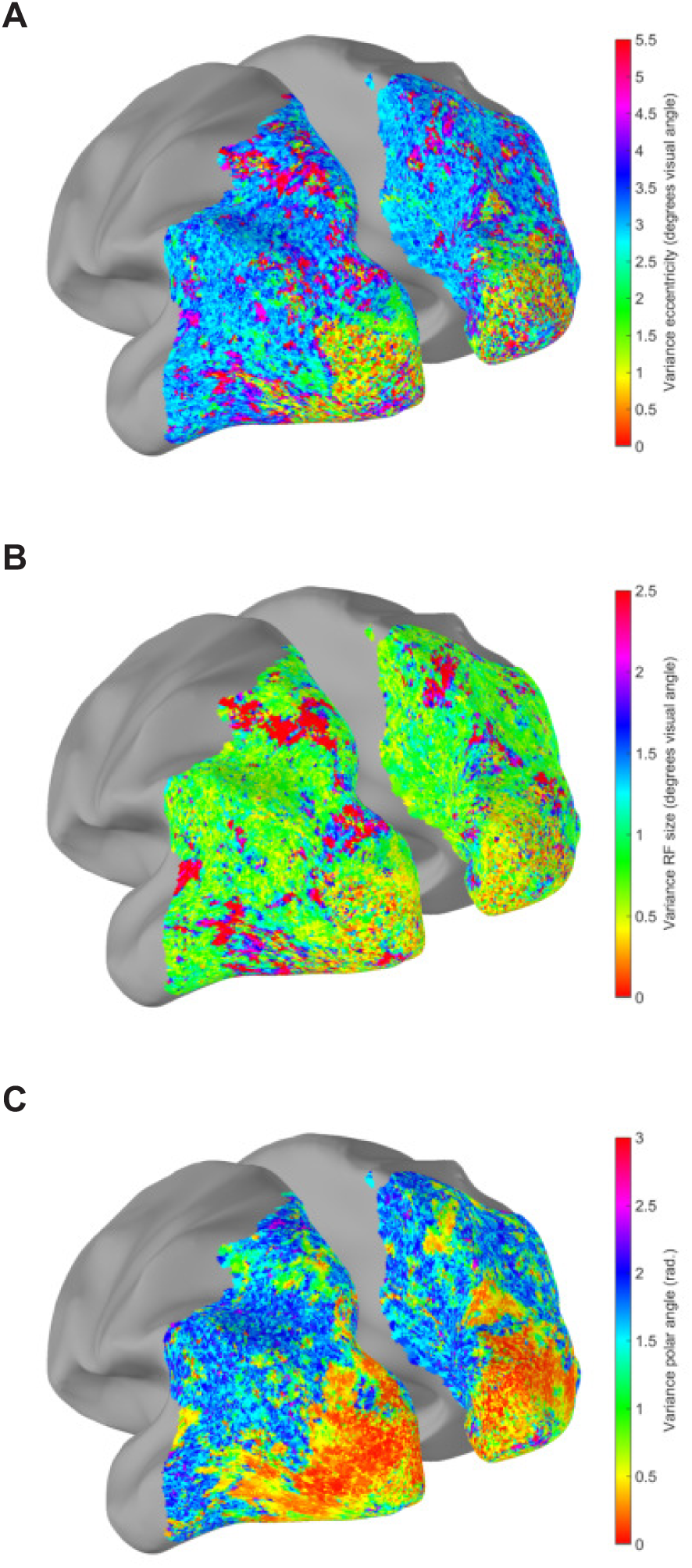
Uncertainty maps of PRFs in the cortex as estimated by our Bayesian approach. From top to bottom: eccentricity, PRF size, polar angle.

Results show clearly that our estimates are better for early visual areas, while becoming worse when we move towards more anterior parts of the brain. The estimates of the measurement noise do not show this difference (Figure 9), which indicates that the Gaussian PRF model is a good model for activities in early visual cortex, but not for activity in higher visual and parietal cortex,consistent with previous findings [11]. The pattern of noise estimates seems to follow the gyri of the brain, which indicates that there might be a physiological feature of folds in the cortex influencing noise levels of voxels.

**Figure 9:**
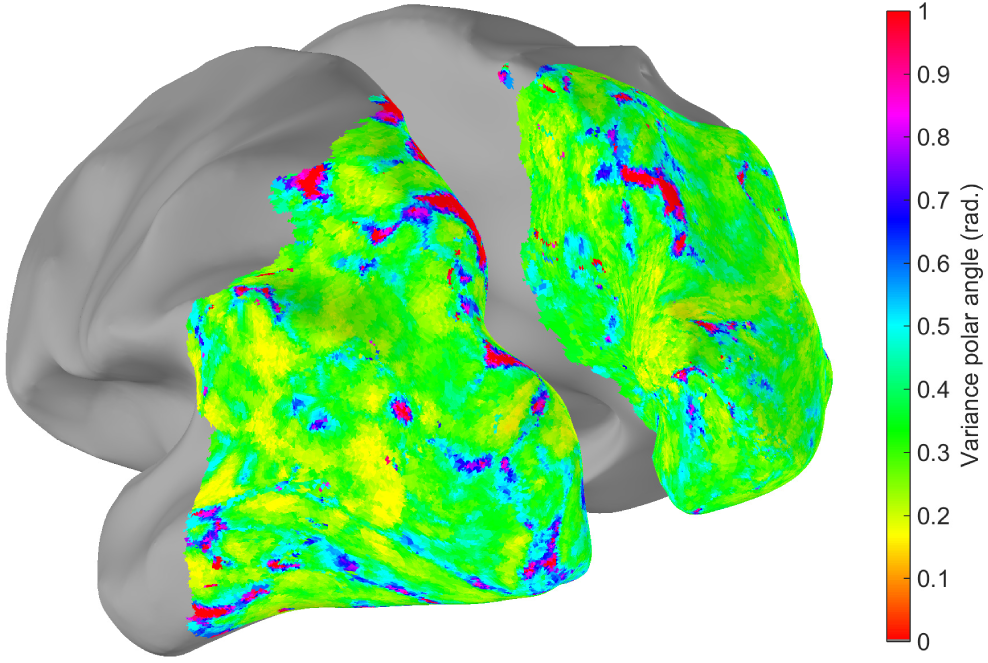
Noise estimation for every voxel.

We compared how much variance our Bayesian estimation explained to the grid search approach and found comparable but slightly lower explained variance with our method (Figure 10). When using a weakly informed prior (see Methods) the explained variance increased slightly, demonstrating that carefully choosing priors could improve our method’s performance.

**Figure 10:**
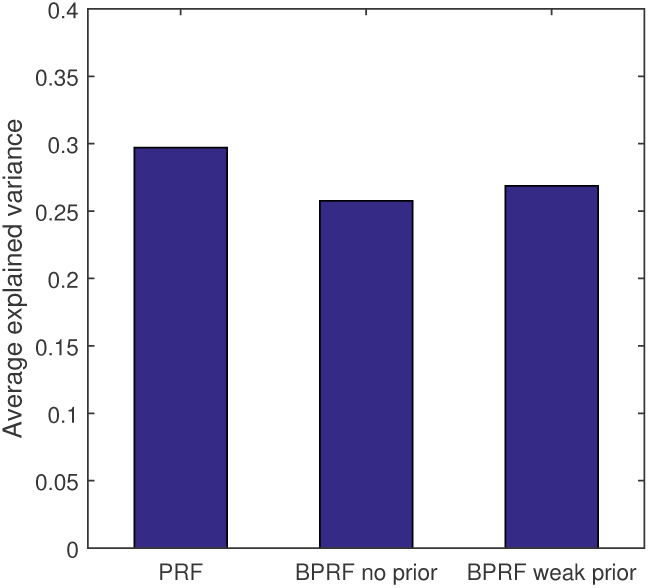
Explained variance for original grid search PRF method and Bayesian PRF method without prior and weak prior.

## 4 Discussion

### 4.1 Improved parameter estimation

Our simulation results show that the Bayesian approach can outperform the conventional PRF method at retrieving the model parameter values that were used in creating our simulated data. On the empirical data our model performed slightly worse then the original grid-search PRF method, but our method provides the added benefit of estimating the full posterior distribution.

### 4.2 Benefits of posterior distribution

A major improvement over the conventional method is that the Bayesian PRF approach adds a measure of uncertainty to each parameter estimate by providing the marginal distributions. First, this allows for direct assessment of how well a model parameter has converged to a single value (even when running a single chain). Indeed, by looking at the variance within each parameters' marginal distribution we can assess the quality of the data. The method also provides an estimate of the noise level, as we sample *σ* a directly using the slice sampler. Secondly, by providing the distribution, this method is able to find interesting information within the distributions. Using the Bayesian approach, one can look at how these distributions behave under different task conditions. For example, interesting attentional effects have been found that imply dynamics in receptive field properties [13, 19]. The distributions might be useful in studying these effects. Furthermore, we can now obtain multimodal marginal distributions. For example, one can imagine a voxel for which two sets of model parameters are equally likely to describe the data. Assuming these parameter values are different for at least one parameter, these would appear as a bimodal distribution in the marginal distributions. Using the conventional PRF method, an underlying bimodal distribution would go unnoticed, as only the best fit in the coarse stage will proceed as a starting point for the fine fit stage.

The Bayesian framework for PRF estimation provides a starting point for future studies. Here we would like to discuss several extensions that can be made to make the framework more versatile.

### 4.3 Choosing priors

In our study we have looked at uninformative as well as informative priors on the parameters of our model. Our weakly informative prior performed slightly better than the uninformative prior. Experiments with strong priors led to heavily biased results that became uninterpretable. Since our priors were based on average relationships over all visual areas, there is significant room of improvement by using area-specific priors. It might also be beneficial to move from an independent prior on receptive field size and position to a combined prior, expressing the relationship between both parameters. Receptive fields near the fovea will then have a higher chance of having small receptive fields where receptive fields in the periphery will have a higher chance of having large receptive fields.

An important extension to the simulation study would be to investigate the effects of boundary conditions on the resulting distributions. Boundary conditions arise when a major part of the population receptive field lies outside the stimulus aperture (the outer radius of the circular stimulus). Due to technical limitations, often the field of view in a MRI scanner is limited to a few degrees of visual angle. As a result, large parts of the receptive field will lie outside this range and only a portion of the receptive field will be used in informing the model. The simulation study could therefore be expanded to analyze effects on the marginal distribution for the parameters on location and size under such boundary conditions. In our simulations we used a stimulus aperture radius of 6.25 degrees visual angle. Therefore, one could look at the performance of retrieving PRF properties for voxels with eccentricities around or exceeding this radius.

### 4.4 Different kernels

Another extension to the Bayesian model would be to use different PRF kernels (i.e. the model for the receptive field). The kernel we used in this simulation study is the two-dimensonal symmetric Gaussian (Equation 7). However, using the conventional PRF method, several kernels have been used [12]. For example, one widely used alternative kernel is the circular symmetric difference-of-Gaussian (DoG), that accounts for surround suppression by modeling a positive center and a negative surround. This and other kinds of kernels can easily be incorporated in the Bayesian framework

### 4.5 Including HRF estimation

In our current approach we used a canonical HRF to map neural activity to BOLD responses. Although the canonical HRF is widely used in FMRI research, there is mounting evidence that the HRF various significantly across voxels [20]. An elegant and likely beneficial extension of the current approach would be to include the estimation of the HRF parameters in the sampling procedure. In addition to our PRF parameters, we can use the data to sample hyperparameters for a parametrized HRF. A widely used parametrization for the canonical HRF is a linear combination of two Gamma functions [21]. Using the Bayesian approach, we can sample the HRF parameters using the slice sampler.

## 5 Conclusion

In conclusion, the Bayesian PRF approach described here provides a new way of estimating PRF properties that builds on the conventional PRF method. The Bayesian method is able to quantify the joint distribution using data and beliefs in parameter behavior from earlier studies, effectively adding a measure of uncertainty to each parameter estimate. Simulations have shown an increased performance of the non-informed Bayesian compared to the conventional PRF method. On empirical data further improvement can be expected by choosing priors more carefully and dealing with boundary effects. The added benefits of our methods will provide a useful tool in unraveling the properties of population receptive fields.

## A Computing the log joint probability density

The log joint PDF of interest is:

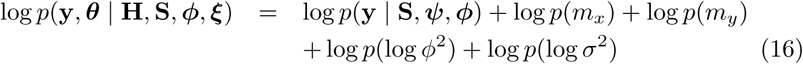

The log PDF of **y** is given by

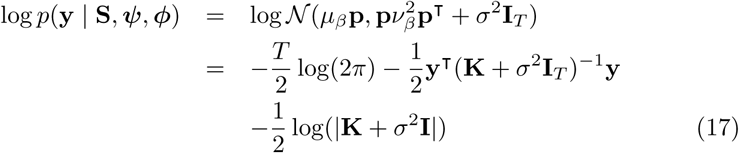

where we used 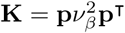. To avoid inversion of a T×T matrix we rewrite:

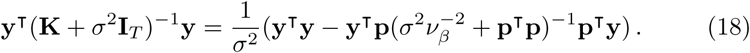

For the determinant we write:

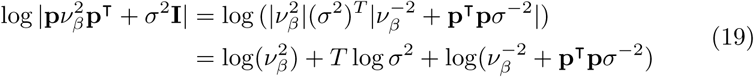

Plugging these into Equation (17) gives:

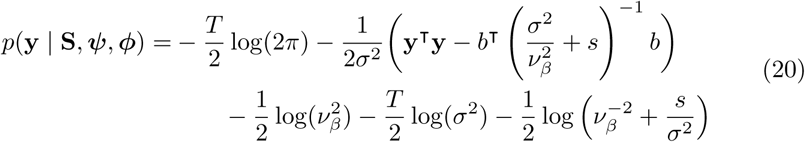

with *b* = **p^T^y**, and *s* = **p^T^p**.

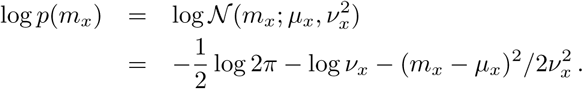

Analogously, we have

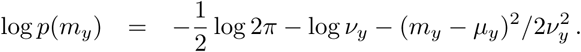

For the kernel width, we have

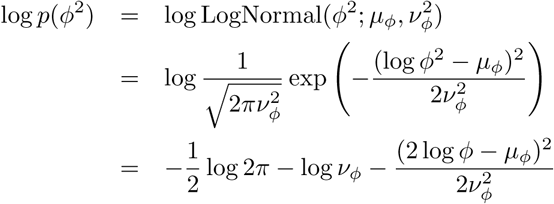

Analogously, a prior on noise could be:

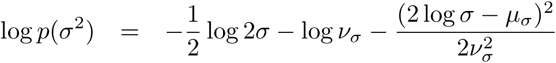

By plugging in the equations while ignoring irrelevant constants, the log joint PDF is proportional to:

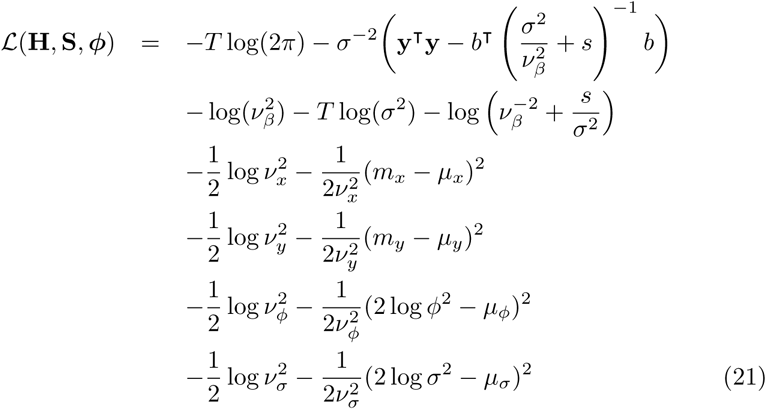

## B Assessing convergence of sampling

To check convergence of our sampling approximation we used the potential scale reduction [22]. By running multiple chains we can check whether our samples mix and dividing our chains in multiple parts enables us to check stationarity. After splitting we have *m* chains of length *n*. For each parameter *ψ* were estimating we compute the within– and between–sequence variance (B and W respectively)

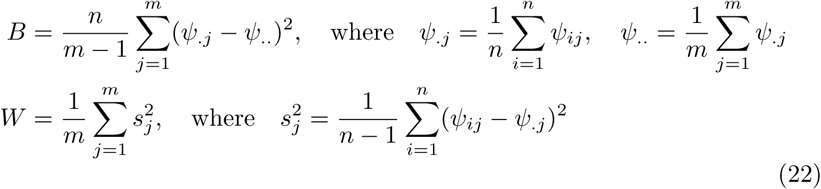

where we label samples as *ψ_i,j_*(*i* = 1,…,*n;j* = 1,…,*m*). From this we can estimate the marginal posterior of the estimated parameter

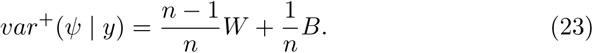

In the limit of *n* → *∞* to, *var*(*ψ* | *y*) approaches *W*. To see how well our estimate has converged for a certain number of samples *n* we compute the potential scale reduction

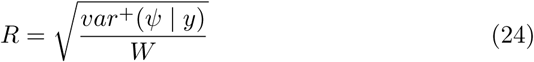

which should converge to 1 for *n* → *∞*. We computed a number of samples such that *R* approached the value of 1 sufficiently.

